# Antibody against envelope protein from human endogenous retrovirus activates neutrophils in systemic lupus erythematosus

**DOI:** 10.1101/776468

**Authors:** Maria Tokuyama, Bronwyn M. Gunn, Arvind Venkataraman, Yong Kong, Insoo Kang, Michael J. Townsend, Karen H. Costenbader, Galit Alter, Akiko Iwasaki

**Author notes:** Corresponding Author: Akiko Iwasaki, Howard Hughes Medical Institute, Department of Immunobiology, Yale School of Medicine, 300 Cedar Street, New Haven, CT 06519, USA.

## Abstract

Neutrophil activation and the formation of neutrophil extracellular trap (NET) are hallmarks of innate immune activation in systemic lupus erythematosus (SLE) and contribute to the systemic interferon signature. Here we report that the expression of an endogenous retrovirus (ERV) locus ERV-K102, encoding an envelope protein, was significantly elevated in SLE patient blood and was correlated with higher interferon status. Induction of ERV-K102 expression most strongly correlated with reduced transcript levels of epigenetic silencing factors. SLE IgG promoted phagocytosis of ERV-K102 envelope protein by neutrophils through immune complex formation. ERV immune complex phagocytosis resulted in subsequent NET formation consisting of DNA, neutrophil elastase, and citrullinated histone H3. Finally, analysis of anti-ERV-K102 IgG in SLE patients showed that IgG2 likely mediates this effect. Together, we identified an immunostimulatory ERV-K envelope protein elevated in SLE that may be a target of SLE IgG and able to promote neutrophil activation.

**eTOC summary:** Using ERVmap, the authors determined that the expression of ERV-K102 locus was elevated in SLE patient blood and correlated with the interferon signature. The envelope protein encoded by this locus activates human neutrophils through immune complex formation with SLE IgG.

## Introduction

Systemic lupus erythematosus (SLE) is a complex and variable autoimmune disease that affects predominantly women of childbearing age. Hallmarks of disease include autoreactive T and B cells, immune complex deposition in tissues, and systemic activation of type I interferon (IFN) signaling and cytokines (Tsokos et al., 2016). Billions of dollars have been spent on research and development and clinical trials over the past few decades, yet belimumab (anti-BAFF monoclonal antibody) is the only FDA-approved targeted therapy for SLE, and it is only effective for roughly half of treated patients (Merrill et al., 2018). Therefore, there is a great need to develop new effective therapies.

Endogenous retroviruses (ERVs) are retroviral sequences that originated from exogenous retroviruses that integrated into our ancestral genome 2 to 40 million years ago and persisted through generations (Stoye, 2012). ERV sequences make up as much as 8% of the human genome, in contrast to the 2% that encodes proteins (Lander et al., 2001). ERVs originally integrated into the genome as proviral sequences, similar to other retroviruses like human immunodeficiency virus (HIV), but most of these sequences have acquired mutations over the course of evolution to render them replication incompetent (Stoye, 2012). In fact, roughly 90% of the ERV sequences are solo-LTRs resulting from homologous recombination between the 5’ and 3’ LTRs that amount to hundreds of thousands of copies in the genome. A minority of ERVs represented in a few thousand copies have a relatively intact proviral structure, comprising of some or all of the original open reading frames (Copeland et al., 1983; Lander et al., 2001).

Solo-LTRs can function as alternative promoters and enhancers are proposed to have contributed to species evolution through regulation of host gene networks and critical host genes, most notably those involved in embryogenesis and stem cell development (Feschotte, 2008; Jern and Coffin, 2008; Chuong et al., 2017; Schlesinger and Goff, 2015; Fuentes et al., 2018). Proviral ERVs have gained growing interest due to their association with diseases such as cancer and neurodegenerative diseases, with particular emphasis on the ERV-K family of ERVs, also known as HML-2 (Schmitt et al., 2013b; Subramanian et al., 2011; Marta et al., 2019). ERV-Ks are the only ERVs that are human-specific with intact open reading frames that remain unfixed in the human population (Jha et al., 2011; Wildschutte et al., 2016; Li et al., 2019). In addition, ERV-Ks are the only ERV family that has been reported to generate viral-like proteins in teratocarcinoma cell line and human blastocysts (Löwer et al., 1993; Bhardwaj et al., 2015; Grow et al., 2015).

ERVs have been implicated in SLE pathogenesis for several decades. Viral antigen related to the primate p30 gag protein have been detected at sites of active lupus glomerulonephritis (Mellors and Mellors, 1976). Antibody reactivity against whole virions, or gag and env peptides from mouse MuLV and baboon BaEV (Blomberg et al., 1994) and ERV-derived ERV-9 and HRES-1 peptides (Blomberg et al., 1994; Bengtsson et al., 1996) have also been observed in SLE. Roughly half of the SLE patients have reactivity against a 28kDa nuclear autoantigen (p28) that is encoded by human T-cell lymphotropic virus (HTLV)-related endogenous sequence (HRES-1) (Banki et al., 1992; Perl et al., 1995). Several haplotypes of HRES-1 contained in the fragile site of chromosome 1 (1q42) are associated with disease (Pullmann et al., 2008). These studies have emphasized the association between ERVs and disease, but there is little understanding of the mechanisms by which they may contribute to systemic inflammation in SLE. Furthermore, the potential roles of other proviral ERV sequences including ERV-K members in SLE have yet to be investigated.

We recently developed a tool called ERVmap to obtain locus-specific proviral ERV transcriptome analysis from RNA sequencing data and revealed over a hundred unique ERV loci significantly elevated in lupus peripheral blood mononuclear cells (Tokuyama et al., 2018). Here we used ERVmap to further analyze an independent cohort of lupus patients to determine the role of proviral ERVs in systemic inflammation and potential mechanisms by which ERVs are dysregulated and contribute to inflammation in SLE.

## Results

### Human-specific envelope-coding ERV-K loci are elevated in lupus blood

Using ERVmap, we observed a global elevation in proviral ERV expression in the whole blood of SLE patients compared with healthy controls in a published RNA sequencing data from a cohort of SLE patients in the rontalizumab in systemic lupus erythematosus (ROSE) trial (Hung et al., 2015). Within this cohort, we identified over 100 significantly elevated ERVs (Figure 1A). The total read counts from elevated ERVs significantly correlated with clinical parameters associated with disease including titers of anti-nuclear antibody (ANA), anti-double stranded DNA (dsDNA), anti-ribonucleoprotein (RNP), and anti-Sm antibodies as well as a decrease in lymphocyte levels and complement C3 levels (Figure 1B). These results show a strong ERV signature that correlate with clinical indicators of SLE.

**Figure 1:**
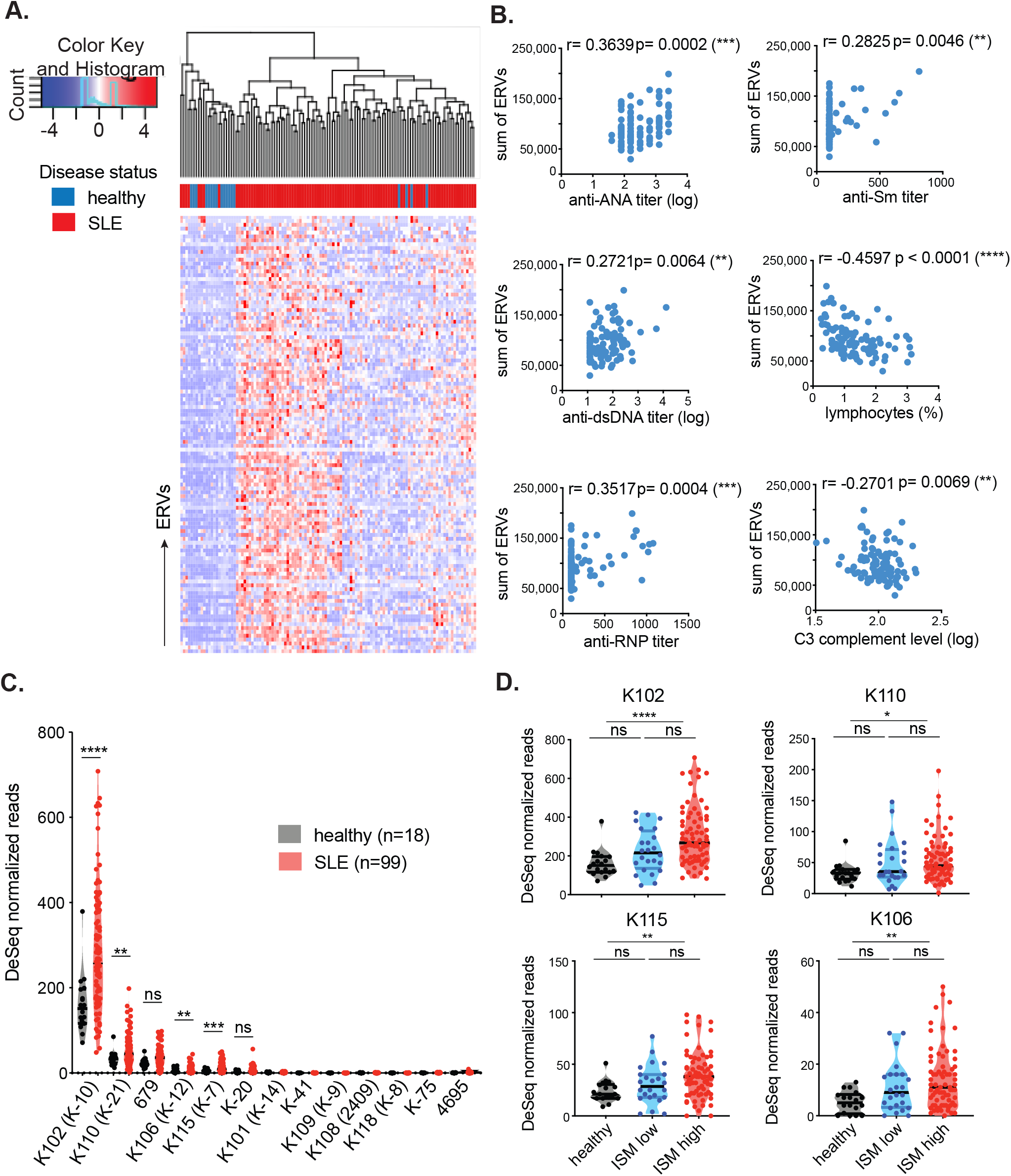
Human-specific envelope-coding ERV-K loci are elevated in lupus blood. ERVmap analysis of RNA sequencing data from whole blood of healthy (n=18) and SLE (n=99) individuals was performed and 113 significantly elevated ERV loci are depicted as a hierarchical cluster map (A). Spearman correlation was calculated between the sum of significantly elevated ERV reads and levels of indicated clinical parameters (B). DESeq normalized read counts for each of the envelope-coding ERV-K loci were compared between healthy and SLE samples; ERVmap IDs are in parentheses (C). Differences between healthy (black), ISM low (blue), and ISM high (red) groups were plotted for the significantly elevated loci (D). Mann-Whitney test was performed to calculate significance for C and D. ISM, IFN signature metric. *, p<0.05; **, p<0.01; ***, p<0.001; ****, p<0.0001; ns, not significant.

ERV-derived envelope (gp70) protein and immune complexes composed of gp70 protein are prevalent in lupus mouse models (Andrews et al., 1978; Izui et al., 1981). In addition, anti-gp70 immune complexes are known to mediate pathology in non-autoimmune mice (Andrews et al., 1978; Izui et al., 1981; Tabata et al., 2000). Based on these findings, we pursued the hypothesis that ERV-derived envelope proteins in humans also have the potential to contribute to SLE and focused specifically on ERV-K (HML-2) members. In the ERVmap database, there are at least 87 ERV-K loci, but only 13 encode a full-length envelope protein without in-frame stop codons (Supplementary Table 1). Based on ERVmap analysis, 4 ERV-K loci with intact coding sequences are significantly elevated in lupus blood compared with healthy controls, K102, K106, K115, and K110 (Figure 1C). In the ERVmap database, these loci correspond to K-10, K-12, K-7, and K-21, respectively, and additional aliases associated with these loci are listed in Supplementary Table 1.

There were significant amino acid sequence homologies between the envelope sequences of the four ERV loci, with up to 97% homology between K102, K115, and K106 and 92% homology between K110 and the other 3 ERV-K loci (Supplementary Figure 1A). The expression levels of these loci correlated within individuals (Supplementary Figure 1B), suggesting that these loci may be coregulated. Based on sequence annotation of these ERVs in the UCSC genome database, K102, K115, K106, and K110 are human-specific ERVs, with no known homology to other primate genomes and do not overlap with other gene loci (Supplementary Figure 1C).

The expression of ERV-K102, K115, K106, and K110 loci were significantly elevated in female patients, but not in male patients, even though the total ERV expression was comparable between the sexes (Supplementary Figure 2A and 2B), suggesting a potential female-bias in the expression of these ERV-K loci. In addition, ERV-K102 expression in particular significantly correlated with the levels of anti-RNP titers, but not with other autoantibody levels (Supplementary Figure 2C). Finally, we observed significant elevation of ERV-K102, K115, K106, and K110 mRNA levels in patients with higher type I interferon (IFN) signature metrics (ISM) (Figure 1D), suggesting a possible role for ERV-K expression in innate immune activation in lupus.

### Transcriptional regulators that correlate with expression of ERV-K102

ERV expression is regulated through epigenetic, transcriptional, and post-transcriptional mechanisms. CpG methylation and H3K9 methylation are key epigenetic modifications that silence the expression of ERVs and are enforced by DNA methyl transferases (DNMTs), histone methyl transferases (HMTs), as well as the nucleosome remodeling and deacetylase (NuRD) complex that removes activating histone acetylation modifications. These modifiers are recruited to specific sites of the genome via sequence specific binding of Kruppel-associated box domain zinc finger proteins (KRAB-ZFPs) and its co-factor KAP1/TRIM28 to transposable elements including ERVs to silence ERV expression upon embryonic development (Ecco et al., 2017). A number of transcription factors are predicted to bind to the LTR of ERVs (Ito et al., 2017). Apolipoprotein B messenger RNA (mRNA)-editing enzyme catalytic polypeptide-like 3 (APOBEC3) family of proteins, tripartite-motif-containing 5a (TRIM5a), and bone marrow stromal cell antigen 2 (BST2, tethrin) are all well-established retroviral restriction factors that restrict other retroviral genomes, including HIV (Malim and Bieniasz, 2012). Given the evidence that these factors also restrict ERV expression in humans and mice (Goff, 2004; Anwar et al., 2013; Ganser-Pornillos and Pornillos, 2019; Treger et al., 2019a; Wolf and Goff, 2008), we sought to determine the potential role of these factors in regulating the expression of ERV-K102, K115, K106, and K110 in lupus samples.

We performed correlation analyses between ERV-K expression and expression of epigenetic silencers, transcription factors known to bind to ERV LTRs, and retroviral restriction factors. A number of epigenetic silencers negatively correlated with ERV expression, including components of the NuRD complex (HDAC1, MTA2, MBD3, CHD4), essential co-factor for KRAB-ZFPs (TRIM28/KAP1), H3K9-specific methyl transferase (ENMT2/G9a), and a protein that interacts with methyl binding proteins and HDAC as a complex (SIN3a) (Figure 2A). ERV-K expression negatively correlated with the transcriptional repressors YY1 and TP53, which have been reported to repress LTR elements (Wang et al., 2007; Wylie et al., 2016), and positively correlated with SP3 and EOMES (Figure 2B). Finally, ERV-K expression did not significantly correlate with retroviral restriction factors in these samples (Figure 2C). The data suggest that ERV-K expression in lupus blood is largely associated with reduced epigenetic and transcriptional silencing in conjunction with transcriptional activation through a few key transcription factors.

**Figure 2:**
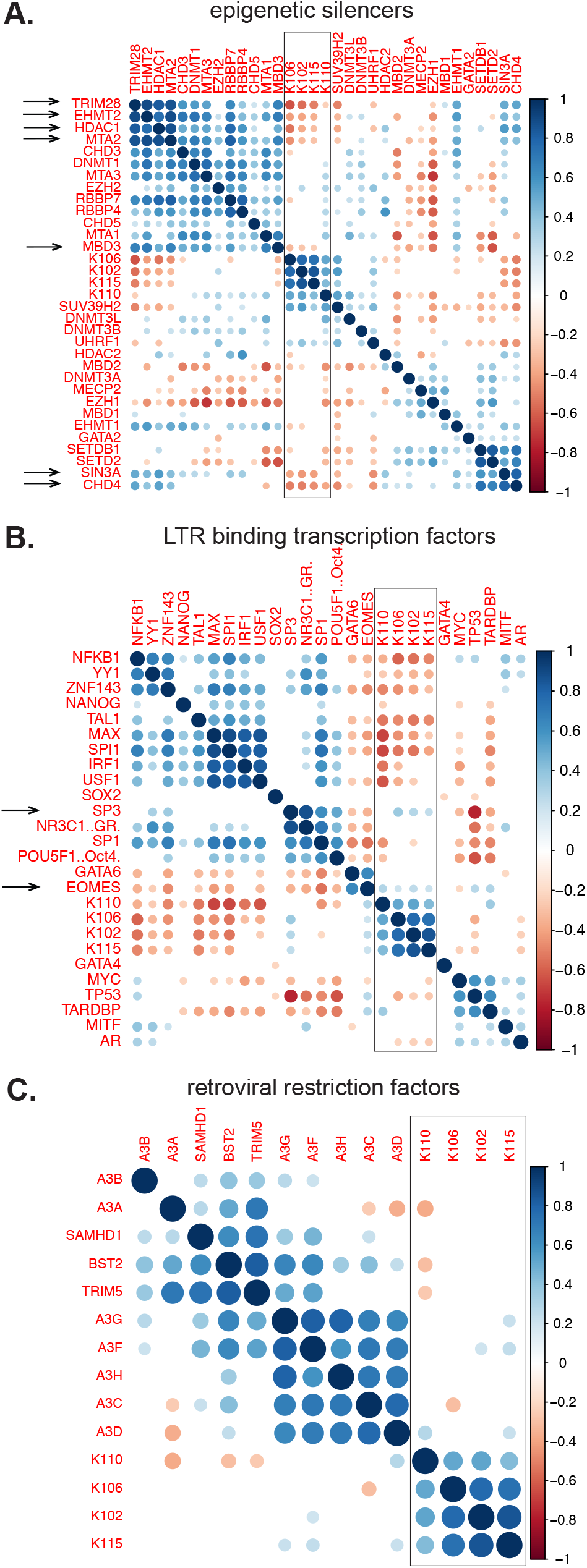
Elevated expression of ERV-K102 most strongly correlates with reduced epigenetic silencing machinery. Using the same RNAseq dataset as the ERVmap analysis, the transcriptome of cellular genes was obtained and expression levels of ERV-K102, K115, K106, and K110 loci were correlated with the expression of epigenetic silencers (A), LTR binding transcription factors (B), and retroviral restriction factors (C) in SLE patient samples (n=99). Correlation plot based on Spearman correlation calculation is depicted. The heatmap colors represent Spearman r values (1 to −1) and only correlations of p<0.05 are displayed. White areas indicate no significant correlation.

### Cloning and generation of recombinant ERV-K102 envelope protein

To determine the exact ERV-K locus predominantly expressed in human PBMCs using a traditional approach, we used a previously described PCR method to amplify the surface unit of ERV-K envelope sequences (Wang-Johanning et al., 2001) from healthy and SLE PBMC RNA (Figure 3A). We observed an expected 1105bp band in both healthy and SLE PBMC samples (Figure 3B). We next cloned the PCR products into a sequencing vector and sequenced multiple colonies per sample. We detected one product from all samples, and blat analysis of this sequence against the hg38 human genome revealed that it is derived from the anti-sense strand of chromosome 1 between 155628270 and 155629354 (1q22), which belongs to the K102 locus, confirming our results from the ERVmap analysis (Figure 3C). Due to the lack of a 292bp deletion observed for type 1 ERV-Ks, we determined that this is a type 2 ERV-K locus that encodes a full-length envelope protein as well as a Rec protein (Ono, 1986; Löwer et al., 1993; 1995). The protein sequence that we obtained was nearly identical to the reference sequence except for two mutations at G208R and T301S in our samples. Based on the latest data available in the 1000 Genomes Project (https://www.internationalgenome.org), no polymorphisms have been reported at these positions. Thus these may be dominant products of somatic mutations in peripheral blood. In order to study the potential role of this envelope protein in disease, we cloned the 1086bp surface unit (SU) of the envelope protein starting at the ATG codon and generated an N-terminal GST-tagged recombinant protein. The production of a 65kDa protein was observed by both Coomassie Blue staining and anti-GST immunoblot of products purified through GST beads (Figure 3D).

**Figure 3:**
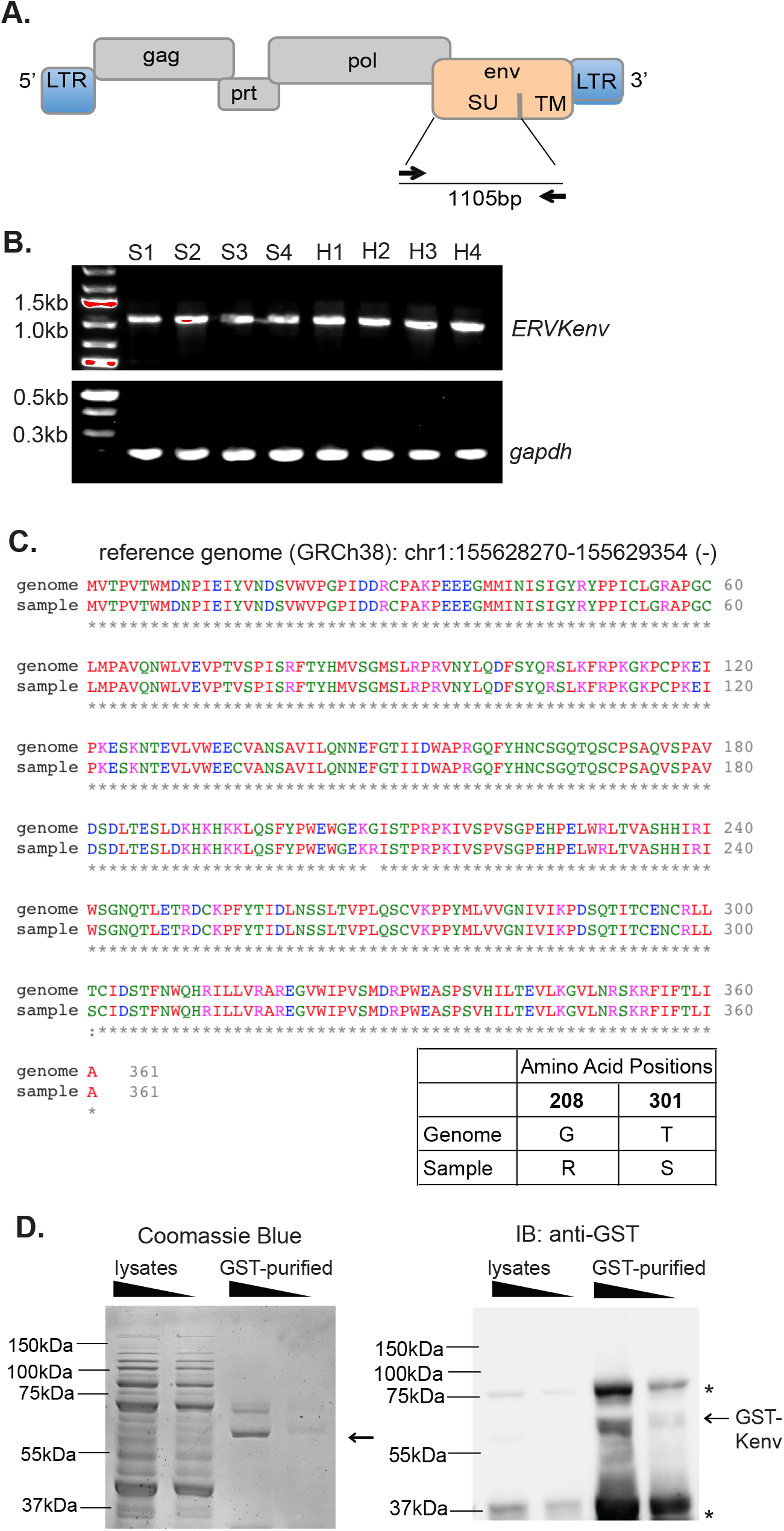
Generation of envelope protein encoded by the ERV-K102 locus. Schematic representation of the proviral structure of ERV-K sequence and the positions for the primers to amplify SU are indicated as arrows (A). Agarose gels for the RT-PCR amplified ERV-K envelope SU and gapdh from healthy (n=4) and SLE (n=4) PBMCs (B). Protein sequence alignment between the reference ERV-K102 sequence (hg38) and the dominant product amplified from PBMCs and the amino acid differences at positions 208 and 301 (C). Coomassie Blue staining and immuno blot with anti-GST antibody for purified ERV-K102 are shown (D). *, non-specific bands.

### IgG from SLE patients activate neutrophils in the form of immune complexes with ERV-K102 envelope

Neutrophils are the most abundant immune cell type in the blood and play a major role in the pathogenesis of lupus disease. Neutrophils are activated by autoantibody immune complexes, and upon activation, secrete intracellular nucleic acids bound by anti-microbial peptides through neutrophil extracellular traps (NETs) composed of autoantigens that perpetuate the IFN response (Lande et al., 2011; Garcia-Romo et al., 2011; Kaplan, 2011; Yu and Su, 2013; Thieblemont et al., 2016). We hypothesized that one mechanism by which ERV-K expression may contribute to the IFN signature is through neutrophil activation by ERV-K immune complexes. To test this, we first measured total IgG against ERV-K102 envelope in healthy and SLE plasma using enzyme-linked immunosorbent assay (ELISA). We observed presence of anti-ERV-K102 IgG in both healthy and SLE plasma at comparable levels (Figure 4A), indicating that there is potential for anti-ERV-K102 IgG in SLE plasma to form immune complexes with ERV-K102 envelope protein. We next tested whether anti-ERV-K102 IgG from SLE patients are capable of inducing ERV-K envelope-specific neutrophil phagocytosis and drive neutrophil activation. We labeled FITC beads with the GST-tagged recombinant ERV-K102 envelope protein, formed immune complexes using either healthy or SLE plasma, and cultured the immune complexes with primary neutrophils isolated from healthy donors. Immune complex phagocytosis was measured by flow cytometry, and antibody-dependent neutrophil phagocytosis (ADNP) score was calculated based on percent FITC+CD3-CD14-CD66+ neutrophils and mean fluorescent intensity (MFI) of FITC (Figure 4B) (Gunn et al., 2018). As a positive control, we generated immune complexes with Ro-SSA, a known autoantigen, and we also generated immune complexes with tetanus and flu HA antigens for comparison. We observed enhanced ADNP of ERV-K102 immune complexes when they were generated with SLE plasma compared with healthy plasma, similar to ADNP levels for Ro-SSA immune complexes (Figure 4C). In contrast, ADNP of tetanus and flu HA immune complexes were comparable between SLE and healthy plasma. In addition, enhanced ADNP of ERV-K immune complexes was observed with purified IgG, indicating that enhanced ADNP by SLE plasma is IgG-mediated (Figure 4D).

**Figure 4:**
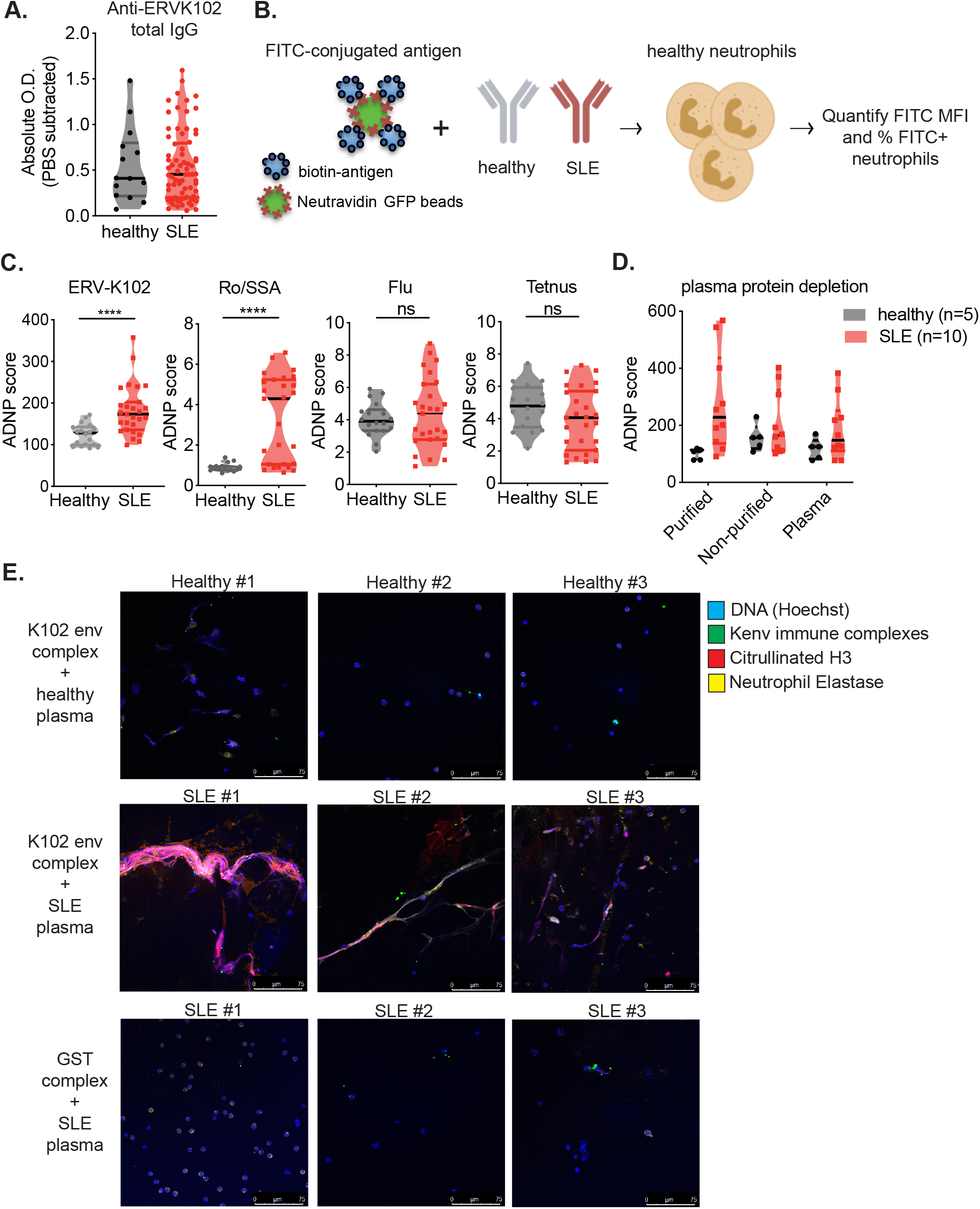
Neutrophil activation by SLE IgG in an immune complex with ERV-K102 envelope protein. Total IgG against ERV-K102 envelope protein was measured by ELISA in healthy (n=14) and SLE (n=73) plasma (A). Neutrophil phagocytosis and NET formation by healthy neutrophils was performed using FITC-conjugated GST-tagged ERV-K102 in an immune complex with either healthy (n=18) or SLE plasma (n=27) (B). ADNP scores were plotted for the indicated antigen immune complexes (C). Mann-Whitney test was performed to calculate significance. ****, p<0.0001; ns, not significant. ADNP with immune complexes generated from IgG-purified healthy or SLE plasma was performed (D). Confocal images of NET formation by activated neutrophils (E). Hoescht (blue), citrullinated histone H3 (red), neutrophil elastase (yellow), and FITC-conjugated immune complexes (green). All experiments are representative of at least two or more repeated experiments.

We next tested whether enhanced neutrophil phagocytosis of ERV-K-immune complex would result in neutrophil activation and thereby NET formation. We incubated healthy neutrophils with ERV-K102 immune complexes generated with either healthy or SLE plasma and stained cells for DNA, citrullinated histone H3, and neutrophil elastase. We observed colocalization of all three markers in a classic NET formation by neutrophils only when neutrophils were stimulated with SLE immune complexes containing ERV-K102 envelope protein, but not with healthy immune complexes or immune complexes generated with just a GST protein (Figure 4E). Together, the data show that in SLE, IgG against ERV-K102 forms immune complexes, which are readily phagocytosed by neutrophils and lead to NET formation.

### Anti-ERV-K102 IgG are predominantly IgG2 and correlates with ADNP

Although levels of total IgG against ERV-K102 envelope protein were comparable between healthy and SLE, we investigated the possibility that reactivity against ERV-K102 envelope protein would differ between IgG subclasses. To determine the antibody profile against ERV-K102 in SLE patients and healthy controls, we performed a subclass-specific binding assay against ERV-K102 as well as known autoantigens such as Ro-SSA, C1q, ssDNA, and collagen. We included tetanus and flu HA antigens for comparison. We observed a subclass-specific response whereby anti-Ro-SSA antibody, as well as those against tetanus and flu HA, were largely targeted by IgG1s. In contrast, anti-ERV-K102 antibody were largely targeted by the IgG2 subclass, but there was no significant difference in levels of anti-ERV-K102 IgG2 between healthy and SLE patients (Figure 5A). ERV-K102 expression also did not correlate with anti-ERV-K102 IgG levels in a smaller cohort of samples (Supplementary Figure 3). We observed segregation in patients that have high levels of anti-Ro-SSA IgG1 and anti-ERV-K102 IgG2. For example, patient #43, 44, 47, 55, 81, and 86 all exhibited high levels of anti-ERV-K102 IgG2 in the absence of high anti-Ro-SSA IgG1 antibody (Figure 5B). These data suggest that patients with higher anti-ERV-K102 IgG2 may define a separate group of patients than those who have high levels of anti-Ro/SSA autoantibodies. Finally, levels of anti-ERV-K102 IgG1 and IgG2 significantly correlated with ADNP of ERV-K immune complexes, suggesting a specific role for anti-ERV-K102 IgG1 and IgG2 in neutrophil activation (Figure 5C).

**Figure 5:**
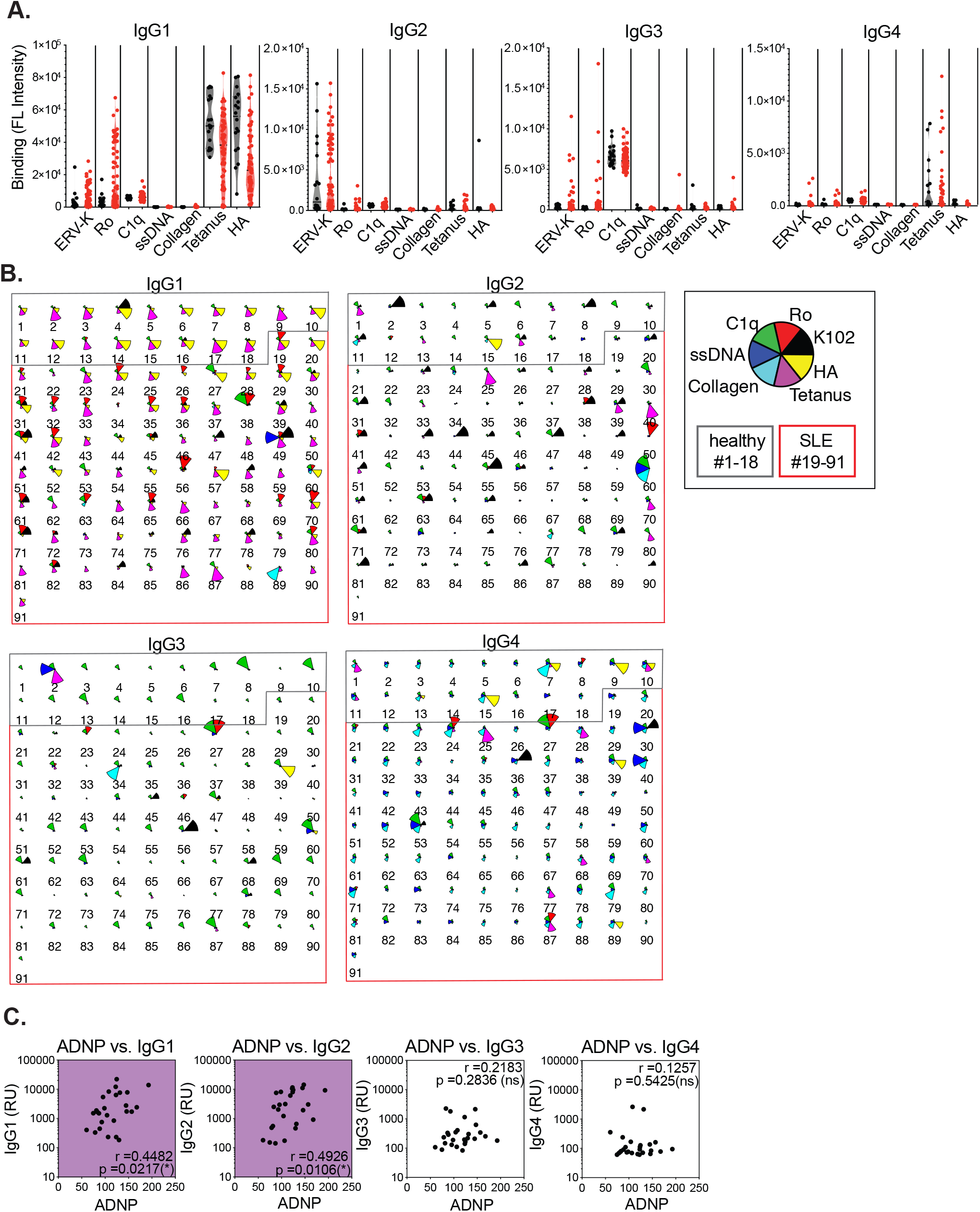
Anti-ERV-K102 envelope IgGs are predominantly IgG2 and correlates with ADNP. Luminex assay was performed on healthy (n=18) and SLE (n=73) plasma against the indicated antigens to determine levels of each of the IgG subclasses (A). Star plot depicting relative levels of IgG against the indicated antigens in different colors showing differences in antibody distribution between individuals (B). Healthy donors (#1-18) in grey box and SLE patients in red box (#19-91). Spearman correlation was calculated between the levels of ADNP and each of the IgG subclasses for SLE patients (n=26). *, p<0.05; ns, not significant.

## Discussion

ERVs have been increasingly associated with diseases ranging from cancer and neurodegenerative diseases to HIV (Rooney et al., 2015; Schmitt et al., 2013a; b; Michaud et al., 2014; Li et al., 2015). In SLE, studies have reported association between ERVs and disease, but they have been limited to a small number of ERV loci (Ogasawara et al., 2000; Fali et al., 2014; Perl et al., 1995). While ERVs have been implicated in lupus pathogenesis (Perl et al., 2009; Nelson et al., 2014; Yu, 2016; Mellors and Mellors, 1976), the mechanisms by which ERVs potentially contribute to systemic inflammation in SLE are still unknown. Given the large abundance of ERV sequences in the human genome, deeper understanding of the functional roles of additional ERV loci may provide better insights into the relationship between ERVs and SLE.

We recently developed a method called ERVmap and found that ERV expression is elevated in SLE patients in our New Haven cohort (Tokuyama et al., 2018). In the present study, we examined a different cohort of SLE patients from the rontalizumab in systemic lupus erythematosus (ROSE) trial (Kalunian et al., 2016) and found that over a hundred ERV loci are elevated in the blood of SLE patients. ERV expression levels showed positive correlation between disease markers such as elevated autoantibodies, decrease in complement proteins, and the IFN signature. Transcriptome analysis further revealed that elevated ERV expression is likely a result of reduced of epigenetic silencing followed by transcriptional activation. We showed that an envelope protein encoded by one of the ERV-K (HML-2) loci is targeted by antibodies generated in SLE and is able to activate neutrophils to secrete NETs following immune complex formation. These data contribute to the greater understanding of the role of ERVs in systemic inflammation and lupus pathogenesis.

Epigenetic changes including DNA methylation and histone modifications are hallmarks of SLE (Ballestar et al., 2006). Drugs that inhibit DNA methylation including hydralazine and procainamide can induce SLE in healthy persons (Ballestar et al., 2006). Based on our current findings that ERV-K expression correlates with reduced epigenetic silencing machinery together with our previous finding of a small subset of KRAB-ZNFs that negatively correlates with ERV elements in SLE patients (Treger et al., 2019b), it is possible that ERV-K expression proceeds reduction in KRAB-ZNF-mediated epigenetic silencing of ERVs. Further experiments will seek to identify the effectors regulating ERV expression in SLE pathogenesis.

Beyond the role of ERV-K102 SU in immune complex-mediated inflammation, there are other potential consequences of elevated expression of this locus. ERV-K102 is one of the hominoid-specific ERVs composed of LTR5_Hs sequence that is predominantly associated with younger ERVs. Although infectious ERVs are considered absent in humans, human tetracarcinoma cell line (Tera-1) produces virions from ERV-K loci including ERV-K102 (Löwer et al., 1993; Bhardwaj et al., 2015). Viral-like particles have also been detected in ERV-K-expressing human blastocytes (Grow et al., 2015) and K102 is associated with virions produced by cord blood mononuclear cells (Laderoute et al., 2007). These data point to the possibility that viral particles containing ERV-K102 viral RNA may arise from specific cell subsets under disease conditions like lupus and mediate virion-associated effects on the host.

Our data showing comparable levels of anti-ERV-K102 IgG in healthy and SLE individuals may imply that ERV envelope is a weak antigen that allows escape of ERV-specific T and B cells from both central and peripheral tolerance mechanisms. Presence of anti-ERV-K102 antibody is therefore a consequence of ERV-K102 and other envelope-coding ERV loci that may become elevated in all individuals under various circumstances. This is consistent with reports showing that antibodies against ERV-K peptides are present in healthy and individuals with autoimmune diseases and ERV env-specific IgG are found in healthy naïve mice at steady state (Herve et al., 2002; Yu et al., 2012).

As the levels of anti-ERV-K102 IgGs were comparable between SLE and healthy controls, it unclear why SLE derived antibodies induced enhanced neutrophil activation. One possibility is that anti-ERV IgG in SLE patients are qualitatively distinct than those found in healthy people. Post-translational modifications of antibodies through glycans dictate effector functions of IgG and differentially impact disease (Gunn and Alter, 2016). In autoimmune diseases, agalactosylated IgG precedes disease onset and reversal of this modification with galactosidases can reverse mouse models of rheumatoid arthritis (Ercan et al., 2010; Ohmi et al., 2016). Therefore, enhanced neutrophil activation by ERV-SLE IgG immune complexes may also be mediated by qualitative differences in SLE IgG, whereby SLE immune complexes differentially engage FcRs on the phagocytes.

Our study demonstrates that ERV envelope is a target of autoantibodies. In lupus disease, such anti-ERV envelope antibody can form immune complexes that are capable of mediating neutrophil activation and NET formation. Given the important role of neutrophils in SLE disease (Kaplan, 2011) and promoting the IFN cascade (Crow, 2014), elevated ERV antigen expression in conjunction with antibody modifications might contribute to the exacerbation of disease. While the underlying cause of SLE remains a mystery, ERVs may be considered as potential autoantigens that stimulate autoantibodies distinct from those against well-established self-antigens.

## Materials and methods

### Patient information

Blood from SLE patients were obtained from two different cohorts. One cohort was recruited from the rheumatology clinic of Yale School of Medicine and Yale New Haven hospital in accordance with a protocol approved by the institutional review committee of Yale University (# 0303025105). The diagnosis of SLE was established according to the 1997 update of the 1982 revised American College of Rheumatology criteria (Hochberg, 1997; TAN et al., 1982). After obtaining informed consent, peripheral blood was collected in EDTA tubes from human subjects and plasma was extracted upon centrifugation. Plasma samples were stored at −80°C. Samples from another cohort was obtained from the SLE Biorepository at Brigham Women’s Hospital. IRB-approved consented whole blood samples were obtained from patients followed in the Brigham and Women’s Hospital Lupus Center (Brigham and Women’s Lupus Center Biobank IRB# 2008P000130). All patients had SLE according to ACR criteria for classification of SLE. Data were collected on age at diagnosis, current age, current SLE disease activity by the SLE disease activity index (SLEDAI)(Lam and Petri, 2005), disease manifestations, past medical history, and past and current medications.

Healthy donor samples were obtained at Yale University School of Medicine in accordance with a protocol approved by the institutional review committee of Yale University (#0409027018). Inclusion criteria for healthy volunteers included age 21-40 or 65 and older, and ability to understand and give informed consent in English. Exclusionary criteria included: current use of medication, such as antibiotics in past two weeks, evidence of acute infection, identified by self-report of fever or symptoms two weeks prior to blood draw, and treatment for cancer in the past three months. At screening (by self-report) women who were pregnant/possibly pregnant were excluded. History of organ, bone marrow or stem cell transplant, liver cirrhosis, kidney disease requiring dialysis, HIV/AIDs, hepatitis C or active hepatitis B, blood donation of 1 pint or more in past 2 months, or treatment with clinical trial medication were also excluded.

Clinical data for SLE patients including baseline levels for ANA, anti-dsDNA, anti-Sm, anti-RNP, anti-La antibodies, lymphocyte counts, and complement levels were obtained as part of the rontalizumab in SLE (ROSE) trial (Kalunian et al., 2015) and shared through an agreement with Genentech.

### RNA sequencing analysis

RNA sequencing data from healthy and SLE whole blood were obtained from a published source (GEO: GSE72509; PRJNA294187) (Hung et al., 2015). Reads were aligned to the human genome (GRCh38), and ERVmap analysis and cellular gene analysis were performed according to previously described methods (Tokuyama et al., 2018). As described, ERV read counts were normalized to size factors obtained through cellular gene analysis, and these normalized counts were used for all subsequent data analysis. Bioconductor R software was used to generate heatmaps, Spearman correlation plots, and star plots.

### Cloning of ERV-K envelope

Peripheral blood mononuclear cells (PBMCs) from Yale healthy donors and SLE patients were obtained through Ficoll-Paque density centrifugation separation. PBMCs were stored in Buffer RLT, and RNA was isolated according to manufacturer’s protocol (RNeasy kit, Qiagen). Reverse-transcription PCR was performed to amplify ERV-K envelope (Wang-Johanning et al., 2001) and *GAPDH* using the following primers:

ERV-K Fwd: 5’ AGAAAAGGGCCTCCACGGAGATG 3’

ERV-K Rev: 5’ ACTGCAATTAAAGTAAAAATGAA 3’

GAPDH Fwd: 5’ CAATGACCCCTTCATTGACC 3’

GAPDH Rev: 5’ GACAAGCTTCCCGTTCTCAG 3’

Amplified products of the expected size for ERV-K envelope were extracted from agarose gels (Zymo Research) and ligated into pCR-Blunt II-TOPO vector for sequencing (Thermo Fisher). Sequencing analysis was performed using ApE software (http://jorgensen.biology.utah.edu/wayned/ape/) and alignment was performed using Clustal Omega (EMBL-EBI). ERV-K envelope protein was cloned out of the sequencing vector and ligated into pGEX-6p-1 N’ GST-tag expression vector (GE Healthcare) between EcoRI and NotI sites using the following primers:

ERV-K EcoRI Fwd: 5’ atcggaattcGTAACACCAGTCACATGGATGG 3’

ERV-K NotI Rev: 5’ atcggcggccgcTGCAATTAAAGTAAAAATGAATCTTTTGGATCTA 3’

### Recombinant protein generation and purification

BL21 strain of *E.coli* were transformed with ERV-K pGEX-6p-1 vector and protein production and GST-bead purification were performed as previously described (Treger et al., 2019b). Transformed cells were grown overnight in YT medium (Sigma-Aldrich) containing ampicillin. Overnight culture was used to inoculate 1L of YT medium and grown until OD_600_ reached 0.6. Cells were cooled in ice-cold water for 10 minutes and grown for 16 to 18 hours in 0.5mM IPTG at 16°C. Cells were pelleted, resuspended in lysis buffer (50mM Tris pH7.4, 100mM NaCl, 0.1% TritonX, 5mM DTT, protease inhibitor complete tablets) at a 1:20 ratio of lysis buffer to starting culture volume, freeze/thawed once, and sonicated in Bioruptor Plus TPX microtubes (Diagenode) for 9 cycles of 30 seconds on and 30 seconds off. Clarified lysates were incubated with Glutathione Sepharose 4B resin (GE Healthcare Life Sciences) at a ratio of 1:40, resin bed volume to lysate volume, for 2 hours at 4°C on a rotator, washed 3 times in PBS containing protease inhibitor, and GST-tagged proteins were eluted 3 times with elution buffer (50mM Tris-HCl pH8.0, 10mM reduced glutathione, protease inhibitor tablets) at a 1:1 ratio of bed volume to elution buffer volume. Eluted proteins were concentrated using Amicon Ultra 0.5ml Centrifugal Filter tubes NMWL 30 KDa (Millipore) and quantified by NanoDrop Spectrophotometer (Thermo Fisher). Lysates and purified products were analyzed by acrylamide gel electrophoresis followed by standard coomassie blue staining and western blot analysis using a rabbit anti-GST Tag polyclonal antibody (CAB4169, Thermo Fisher).

### Immune complex generation

Immune complexes were generated using human plasma and recombinant protein as previously described (Gunn et al., 2018). Ro-SSA antigen was purchased (Arotec Diagnostics) and HA and tetanus proteins were obtained from ImmuneTechnology Corp. and MasBiologics, respectively. Recombinant proteins were biotinylated with EZ-Link Sulfo NHS-LC-LC biotin (Thermo Fisher) at a 50 molar excess for 30 minutes at room temperature. Excess biotin was removed using 7K MWCO, Zeba Spin Desalting Columns (Thermo Fisher) according to manufacturer’s protocol. Biotinylated proteins were coupled to FITC-labeled 1um FluoSpheres NeutrAvidin-labeled Microspheres (Thermo Fisher) at 1 ug to 1 ul ratio of protein to beads for 2 hours at 37°C, washed twice in 0.1% BSA in PBS, and resuspended in 1ml of 0.1% BSA in PBS for 10ug of protein. 0.1ug of bead-coupled proteins were incubated with 100ul of 1:100 dilution of plasma IgG in a 96-well plate for 2 hours at 37°C to generate immune complexes, and beads were pelleted by centrifugation at 2000rpm for 10 minutes. For IgG purification, plasma was diluted 1:10 in Melon Gel Purification Buffer and Melon Gel IgG Spin Purification Kit (Thermo Fisher) was used according to manufacturer’s protocol. Purified IgG was used at a final concentration of 1:100 to generate immune complexes as above.

### Neutrophil phagocytosis

Healthy polymorphonuclear neutrophils (PMNs) were obtained by treating healthy whole blood with ACK lysis buffer (Thermo Fisher) for 5 minutes at room temperature, followed by centrifugation and a PBS wash. Cells were resuspended in RPMI medium (10% FBS, 1% P/S, HEPES, L-glut), and 50K cells were incubated per well of immune complexes generated as described above in 200ul for 1 hour at 37°C. Cells were pelleted and stained for CD66b, CD14, and CD3 and analyzed by BD LSRII flow cytometer to quantify mean fluorescent intensity (MFI) and percentage of CD3-CD14-CD66b+FITC+ cells. ADNP score was calculated by (FITC MFI) x (% FITC+) / 10,000 for each well and the average of duplicate wells was obtained.

### Microscopy

For NET analysis, 40K PMNs were plated on poly-L-lysine coverslips in a 24 well plate for 15 minutes at 37°C and unbound cells were washed off with PBS. Each well was incubated with 500ul of 0.1ug immune complex as described above for 2 to 3 hours at 37°C. Cells were fixed with 4% paraformaldehyde for 15 minutes at room temperature, washed in PBS, and blocked overnight in 2mM EDTA PBS containing 10% FBS, 1% BSA, 0.05% Tween 20 at 4°C. Cells were sequentially stained with the following antibodies: mouse anti-neutrophil elastase (MABS461, Millipore) at 1:250, Cy3 anti-mouse IgG (Jackson ImmunoResearch) at 1:1000, rabbit anti-histone H3 (Ab5103, Abcam) at 1:250, and Cy5 anti-rabbit IgG (Jackson ImmunoResearch) at 1:1000. Cells were then stained with Hoechst 33342 (Thermo Fisher) at 1:100 for 10 minutes at room temperature and ProLong Gold Antifade Mountant (Thermo Fisher) was added along with cover slips. Leica TCS SP8 confocal microscope using the 40x immersion lens was used to obtain images of NETs.

### Antibody profiling analysis

Antibody subclass profiling analysis was performed as previously described (Brown et al., 2012). Briefly, recombinant K102, C1q, ssDNA, Ro-SSA, collagen, HA, or tetanus toxin were coupled to MagPlex beads (Luminex) via sulfo-NHS coupling chemistry. Samples were diluted 1:1000 (IgG1) or 1:100 (IgG2, IgG3, IgG4) in 1X PBS + 0.1% bovine serum albumin (BSA) + 0.05% Tween20 and incubated with antigen-coupled beads for 2 hours at room temperature with shaking. Beads were washed, and different antibody subclasses (IgG1, IgG2, IgG3, IgG4) were detected by incubating with 0.65µg/ml of PE-labeled secondary antibodies (Southern Biotech) for 1 hour at room temperature with shaking. Beads were washed and analyzed on a Flexmap 3D instrument (Luminex). The median fluorescent intensity of 30 beads/region was recorded.

### Statistical analysis

Graph Pad Prism (v8.0) was used for all statistical analysis. Non-parametric Mann-Whitney t-test was performed to calculate significance between groups. To compare more than two groups, we used one-way ANOVA Kruskal-Wallis test. Spearman r was calculated to determine significant correlation. Data are represented as means ± SEM. In all cases, *, p<0.05; **, p<0.01; ***, p<0.001; ****, p<0.0001; ns, not significant.

## Supplementary materials

Supplementary Figure 1: Sequence analysis of elevated ERV-K loci.

Supplementary Figure 2: ERV-K102 expression based on sex and anti-RNP IgG level.

Supplementary Figure 3: Correlation between ERV-K102 and anti-ERV-K102 IgG levels.

Supplementary Table 1: List of envelope-coding ERV-K loci in ERVmap database.

## Author contributions

M. Tokuyama and A. Iwasaki conceptualized the study, designed experiments, and interpreted all the data. M. Tokuyama and A. Iwasaki wrote the manuscript with input from co-authors. M. Tokuyama performed all experiments, except antibody profiling experiment performed by B. M. Gunn. B. M. Gunn and G. Alter were instrumental in ADNP experiments and provided necessary reagents. A. Venkataraman performed ERV-K PCR and generated ERV-K envelope protein with the guidance of M. Tokuyama. Y. Kong performed raw RNA sequencing analysis. I. Kang and K. Costenbader provided SLE patient samples and provided clinical research guidance. M. Townsend provided clinical data from ROSE trial and provided clinical research guidance. A. Iwasaki supervised and acquired funding for the study.

## Acknowledgement

We thank Allison Nelson, clinical research nurse (Yale University), Albert Shaw, M.D., Ph.D. (Yale University), and Ruth Montgomery, Ph.D. (Yale University) for help with sample acquisition. This work was supported in part by grants provided by AbbVie (to A.I.), Translation Accelerator Grant from the Brigham and Women’s Hospital (to A.I.), and NIH awards T32 AI89704-3 (to M.T.). A.I. is an investigator of the Howard Hughes Medical Institute. K. Costenbader is supported by NIH R01 AR057327 and K24 AR066109. G. Alter is supported by the Samana Kay MGH Scholar Award and the Ragon Institute.

