## Supplementary figures and table for "Antibody against envelope protein from human endogenous retrovirus activates neutrophils in systemic lupus erythematosus"

Suppl Figure 1. Sequence analysis of elevated ERVKs

A.

| | K110 | K115 | K106 | K102 |
|---|---|---|---|---|
| **K110** | 100 | 92.54 | 93.92 | 93.92 |
| **K115** | 92.54 | 100 | 96.96 | 97.51 |
| **K106** | 93.92 | 96.96 | 100 | 97.79 |
| **K102** | 93.92 | 97.51 | 97.97 | 100 |

B.

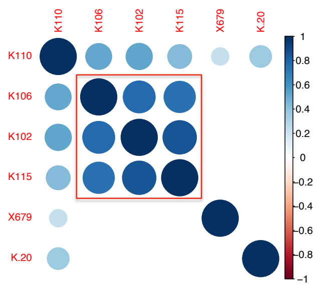


C.

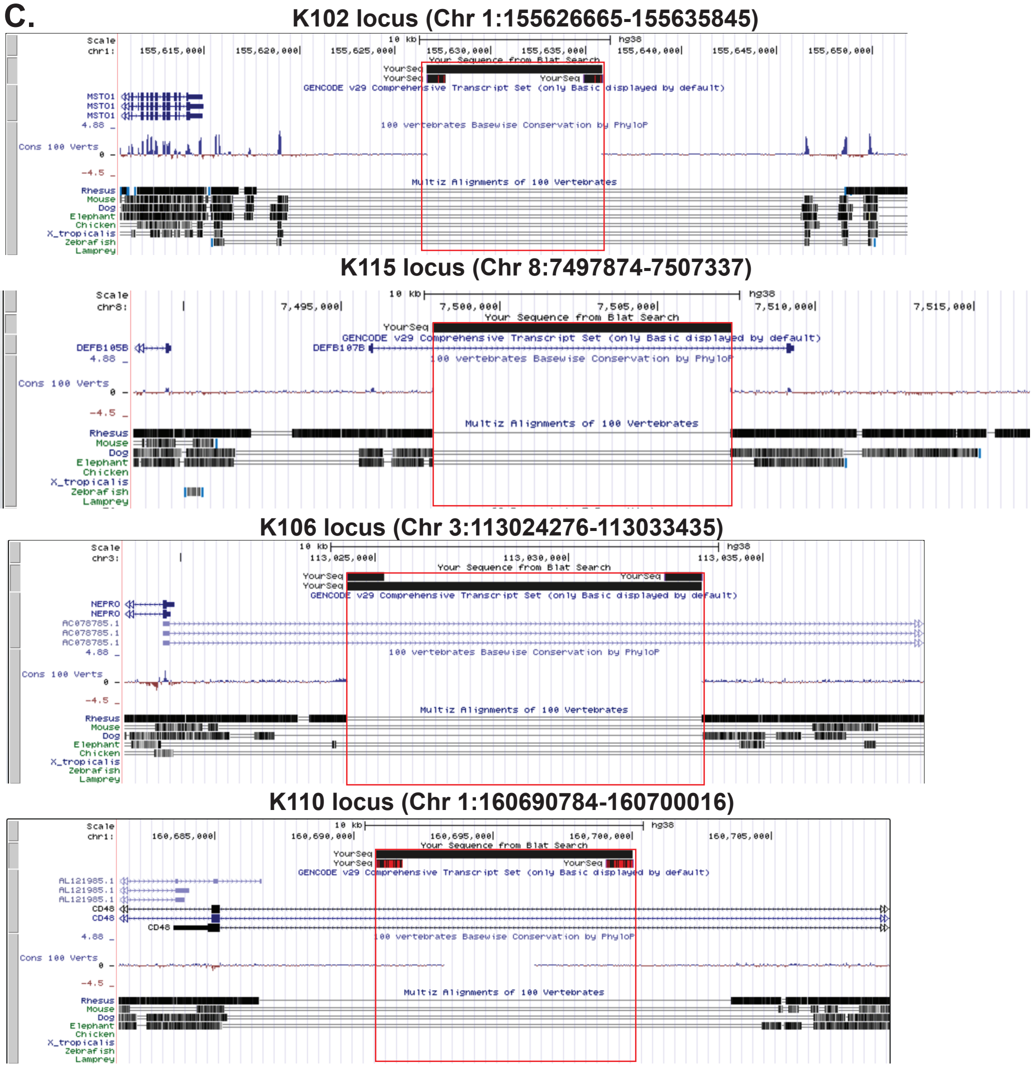


**Supplementary Figure 1: ERV-K102, K115, K106, and K110 loci are human-specific and envelope SU are highly similar in sequence.** Percent homology between the envelope SU sequences of the 4 ERV-K loci at the amino acid level are indicated in a matrix (A). Spearman correlation was calculated between expression levels of these ERV-K loci in SLE samples (n=99) and plotted as a correlation plot (B). Red box indicates high correlation between the indicated loci. UCSC genome browser outputs are displayed for each of the locus to show absence of sequence conservation in primates and other species (C). The black box at the top indicate the ERV-K locus.

Supplementary Figure 2

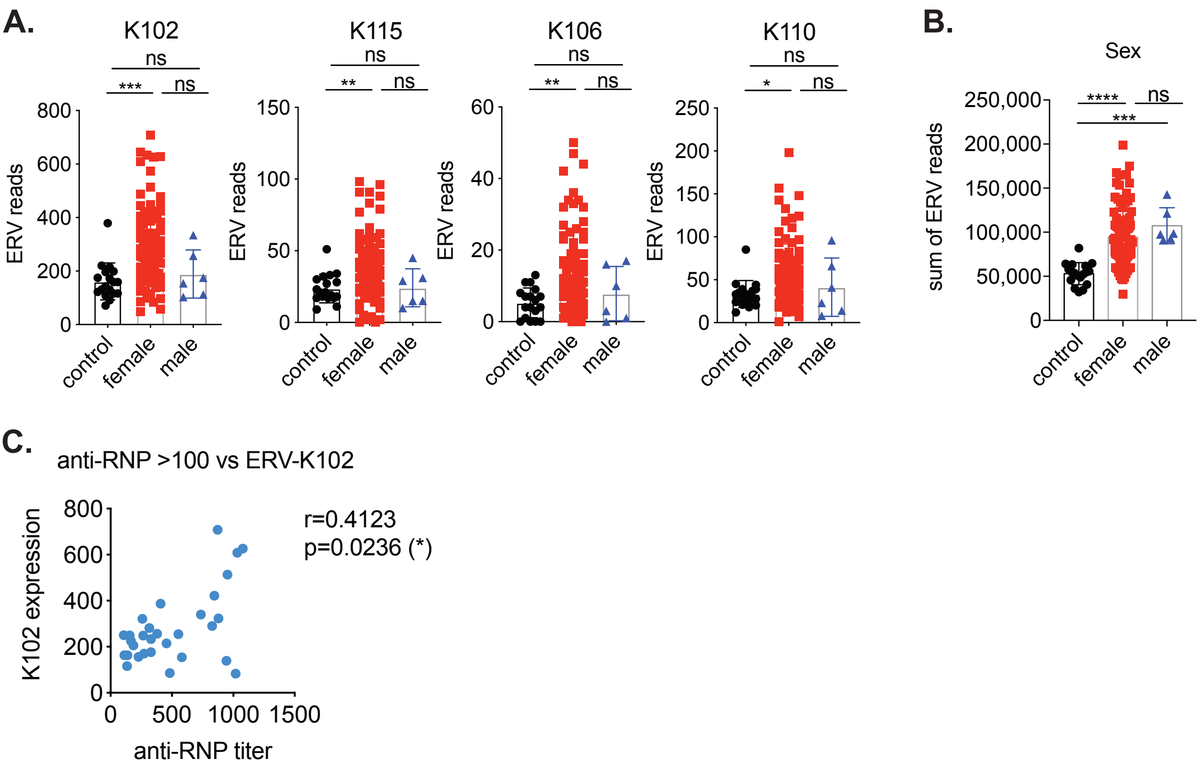


**Supplementary Figure 2: ERV-K expression is higher in females than males and correlates with anti-RNP titer.** Expression levels of the indicated ERV-K loci (A) or sum of reads from the significantly elevated ERVs (B) were differentially plotted depending on the sex of the individual (control, n=18; females, n=93; males, n=6). Non-parametric one-way ANOVA analysis was performed to calculate statistical significance between each groups. C) Correlation plot of ERV-K-10 (K102) expression level and titers of anti-RNP only for SLE patients that anti-RNP antibody titers over 100 (n= 30). *, $p<0.05$; **, $p<0.01$; ***, $p<0.001$; ****, $p<0.0001$; ns, not significant.

Supplementary Figure 3

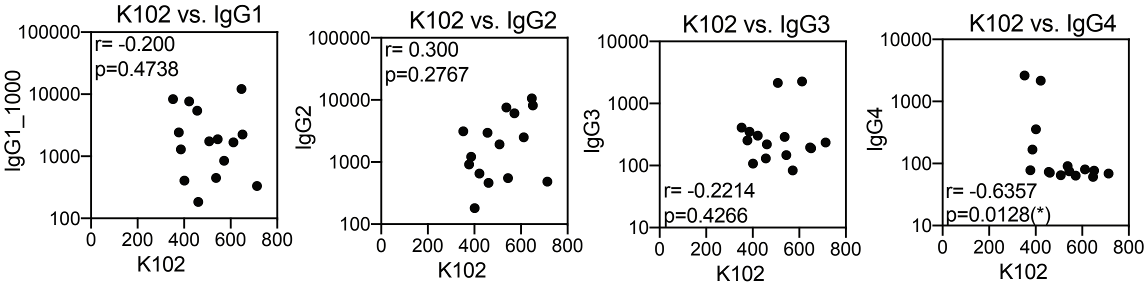


**Supplementary Figure 3: No correlation between ERV-K102 expression and anti-ERV-K102 IgG.** Correlation plots of ERV-K102 expression and the titers of anti-ERV-K102 IgG for each of the isotypes in SLE patients (n=15). Spearman correlation was calculated and both the rho and p values are indicated for each correlation plot.

Supplementary Table 1

**Intact envelope ORF**

| Chr | Start | End | ERVmap ID | Aliases |
|---|---|---|---|---|
| 1 | 155626665 | 155635845 | K-10** | K102, K(C1b), K50a, ERVK-7 |
| 1 | 160690784 | 160700016 | K-21** | K110, K(C1a), ERVK-18, K18 |
| 2 | 129961897 | 129969685 | 679‡ | K-76**, K(2q21.1) |
| 3 | 113024276 | 113033435 | K-12** | K106, K68, ERVK-3, K(C3) |
| 6 | 42893670 | 42903629 | K-41** | K(OLDAL035587),KOLD35587 |
| 6 | 77716944 | 77726366 | K-9** | K109, ERVK-9, K(C6) |
| 7 | 4590930 | 4600359 | 2409‡ | K108, ERVK-6 |
| 8 | 7497874 | 7507337 | K-7** | K115, ERVK-8 |
| 11 | 118721014 | 118730174 | K-20** | ERVK-20, K(C11b),K37 |
| 11 | 101695062 | 101704528 | K-8** | K118,ERVK-25, K(C11c), K36 |
| 16 | 34997025 | 34999771 | K-75** | K(19p13.3) |
| 19 | 27637114 | 27646451 | 4695‡ | HERVK-28, K(C19), ERVK-19 |
| 22 | 18938673 | 18947848 | K-14** | K101, K(C22), ERVK-24 |

**No intact envelope ORF**

| Chr | Start | End | ERVmap ID | Aliases |
|---|---|---|---|---|
| 1 | 207632285 | 207641252 | 6171‡ | K-79**, K(1q32.2) |
| 1 | 238762294 | 238764473 | K-34** | K(4p16.3b) |
| 1 | 166605365 | 166611021 | K-66** | K12 |
| 2 | 186812953 | 186813479 | HERVK30 | |
| 3 | 101692114 | 101701253 | 1045‡ | K-18**, K(II), ERVK-5 |
| 3 | 148552182 | 148567777 | 1163‡ | K-52**, ERVK-13 |
| 3 | 9841750 | 9854710 | 864‡ | K-53**, K11, ERVK-2 |
| 3 | 75551278 | 75559986 | 985‡ | K-39**, K(3p12.3) |
| 3 | 185562547 | 185571727 | K-15** | K117, ERVK-11, K50b |
| 3 | 125890458 | 125899596 | K-19** | K(I), ERVK-4 |
| 4 | 240028 | 246321 | 1284‡ | K-82**, K(4p16.3a) |
| 4 | 9121945 | 9131359 | 1303‡ | K17b |
| 4 | 164995690 | 165005061 | 1685‡ | K-55**, K5, ERVK-12 |
| 4 | 190106257 | 190113553 | 1741‡ | K(5p13.3) |
| 4 | 3977323 | 3986912 | K-35** | K77 |
| 4 | 9657955 | 9667550 | K-36** | K50c |
| 4 | 68597990 | 68603505 | K-63** | K(4q32.1) |
| 4 | 160658785 | 160661286 | K-83** | K(4q32.3) |
| 5 | 30485109 | 30495950 | 1793‡ | K-16**, K104,K50d |
| 5 | 71571927 | 71578401 | 1878‡ | ERVK3-3, ML6-c5 |
| 5 | 93431619 | 93432157 | HERVK31 | |
| 5 | 156657705 | 156666885 | K-11** | K107, K(C5), ERVK-10 |
| 5 | 46000056 | 46009900 | K-28** | K(5q33.3) |
| 5 | 154635952 | 154644655 | K-42** | K18b |
| 5 | 136413881 | 136414650 | K-81** | |
| 6 | 60652252 | 60663180 | 2153‡ | K-59**, K23 |
| 6 | 150856845 | 150862887 | 2373‡ | K-72**, K(7p22.1a) |
| 6 | 28682589 | 28692958 | K-29** | K(OLDAL121932), K69, K20 |
| 6 | 164023 | 174392 | K-30** | |
| 6 | 3054799 | 3055508 | K-78** | |
| 7 | 104747921 | 104752819 | K-47** | ERVK-14 |
| 7 | 141751125 | 141756138 | K-48** | ERVK-15, K(OLDAC004979) |

| | | | | |
|---|---|---|---|---|
| 8 | 12216469 | 12225829 | 2734‡ | K(8p23.1d) |
| 8 | 17907211 | 17916624 | 2744‡ | K(8q24.3b) |
| 8 | 139459107 | 139470560 | 2976‡ | K-74**, K(9q34.11) |
| 8 | 145021244 | 145028842 | 2985‡ | K-61**, K29 |
| 8 | 12458983 | 12468498 | HERVK-26 | KOLD130352 |
| 8 | 8197177 | 8206699 | K-38** | K27 |
| 8 | 46264027 | 46272039 | K-62** | K70,K43 |
| 9 | 136780314 | 136789786 | 3177‡ | K-43**, K30 |
| 9 | 128850235 | 128857457 | K-60** | K31 |
| 10 | 99818702 | 99827957 | 3340‡ | ERVK-17, K-24**, c10_B |
| 10 | 6824178 | 6833641 | K-22** | ERVK-16, K(C11a),K33 |
| 10 | 3393039 | 3402593 | K-50** | |
| 11 | 3447426 | 3456906 | 3379‡ | K7 |
| 11 | 7898106 | 7906049 | 3400‡ | ERVK3-4 |
| 11 | 58996951 | 59006462 | 3505‡ | K-67**, K(11q12.3) |
| 11 | 62368490 | 62383091 | HERVK-27 | K(OLDAC004127) |
| 11 | 71764073 | 71767231 | K-51** | |
| 12 | 111816238 | 111825646 | 3902‡ | ERVK3-5 |
| 12 | 133090536 | 133096478 | 3922‡ | K42 |
| 12 | 58327458 | 58336915 | K-13** | K119, ERVK-21, K(C12), K41 |
| 12 | 34619619 | 34629282 | K-44** | K50e |
| 12 | 110570037 | 110571520 | K-69** | K(16p11.2) |
| 14 | 24009540 | 24015782 | 4086‡ | K(OLDAL136419), K71 |
| 14 | 105673312 | 105676203 | K-68** | K(15q25.2) |
| 17 | 8056336 | 8063901 | K-49** | K(19q13.41) |
| 19 | 22575531 | 22583732 | 4669‡ | K-54**, K51 |
| 19 | 52740546 | 52749825 | 4768‡ | K-58**, K(22q11.23) |
| 19 | 53359490 | 53365214 | 4776‡ | K-73**, LTR13 |
| 19 | 37106647 | 37116164 | HERVK-29 | K(OLDAC012309),KOLD12309 |
| 19 | 20276590 | 20286703 | K-26** | K52 |
| 19 | 35572304 | 35576532 | K-57** | K(19q13.12b) |
| 19 | 385098 | 387637 | K-77** | ERVK-22 |
| 20 | 34126943 | 34136578 | K-46** | K(OLDAL136419),K59 |
| 21 | 18561004 | 18570006 | 4873‡ | K-23**, K60, ERVK-23 |
| 22 | 23538316 | 23548383 | 6272‡ | KOLD345, K(OLDAP000345) |
| X | 62740126 | 62747923 | 5423‡ | K-71**, K(Xq11.1) |
| X | 66460972 | 66470510 | 5449‡ | K-40**, K(Xq12) |
| X | 154589363 | 154594514 | 5768‡ | K-65**, K63 |
| X | 154608422 | 154615762 | K-64** | K63 |
| Y | 6958031 | 6965791 | 5042‡ | K-70**, K(1q43) |
| Y | 24252613 | 24255152 | 5252‡ | K(Yq11.23a) |
| Y | 25412809 | 25417531 | 5269‡ | K(Yq11.23b) |

* Originally named in Mayer J. et al. Mobile DNA 2011, but updated in Schmitt K. et al. JVI 2013.

** ERV-K loci listed in Schmitt K. et al. GBE 2013. IDs were assigned in order annotated.

† ERV-W loci listed in Schmitt K. et al. JVI 2013. IDs were assigned in order annotated.

‡ Numerical identifier for ERVs are the same as annotated in Vargiu L. et al. Retrovirology 2016.
